# Flexible oocyte manipulation with delayed maturation and improved SCNT efficiency using induced pluripotent stem cells

**DOI:** 10.1101/2025.10.14.682316

**Authors:** Luis H. de Aguiar, Yoke Lee Lee, Abdallah W. Abdelhady, Prasanthi Koganti, Vimal Selvaraj, Soon Hon Cheong

## Abstract

Somatic cell nuclear transfer (SCNT) remains inefficient, limiting its practical use in cattle reproduction and research. This study investigated two complementary strategies to enhance handmade cloning (HMC): (1) holding bovine oocytes overnight and delaying maturation to enable a second round of SCNT and (2) using bovine-induced pluripotent stem cells (biPSCs) as donor nuclei to enhance developmental competence. Bovine oocytes were subjected to either conventional *in vitro* maturation (CONV; 20 h) or delayed maturation using a holding medium for 20 h before CONV (HOLD). Matured oocytes were used for SCNT, parthenogenetic activation (PA), or *in vitro* fertilization (IVF) as controls. Handmade SCNT embryos were reconstructed using fibroblasts or biPSCs as donors, activated, and cultured for 7 days. Results showed no significant differences between CONV and HOLD groups in oocyte maturation, recovery after stripping, survival after zona removal, or cleavage and blastocyst development after SCNT. Fusion rates using fibroblasts were comparable between groups (42.6±6.0% vs. 50.3±9.8%), with biPSCs showing significantly higher fusion rates in CONV group (85.7±8.2% vs. 50.5±8.8%, P<0.05). Among fused embryos, biPSCs produced higher blastocyst rates (33.3±16.7%) compared with fibroblast donors (21.9±12.6%, P<0.05). Across all reconstructed embryos, cleavage and blastocyst development were also greater with biPSCs (odds ratios 3.4 and 2.7 respectively). These findings indicate that delaying maturation offers flexible timing for SCNT without compromising competence. Moreover, biPSCs enhance embryo developmental outcomes, supporting their use as superior donor cells for advancing cloning efficiency and applications in reproductive biotechnology.

## 1. Introduction

Somatic cell nuclear transfer (SCNT) is a powerful technique that enables the generation of viable offspring from preserved somatic cells, with applications spanning agriculture, companion animals, wildlife conservation, and biotechnology. Since the birth of the first calf via SCNT [1], numerous refinements have been made to improve efficiency; however, the fundamental principles of the technique remain unchanged. Traditional SCNT involves the injection and fusion of a donor cell with an enucleated mature oocyte using micromanipulation [2], followed by activation and embryo culture. Handmade cloning (HMC) is a modified version of this method [3], that eliminates the need for micromanipulators. However, SCNT efficiency continues to be low, where only 1-5% of the embryos transferred produce a live birth [4]. Therefore, hundreds of oocytes have to be manipulated individually to obtain a single cloned animal. For HMC, even more oocytes might be needed since up to 50% of the cytoplasm is discarded during bisection and enucleation [5]. The procedure for SCNT is highly technical and time-consuming which limits the number of oocytes that can be processed within a specific time frame. Furthermore, to minimize oocyte aging and decrease embryo development, activation is typically performed before 30 hours post-maturation [6], restricting the window for oocyte manipulation to a single working day. Extending the usability of oocytes beyond one day could increase the total SCNT embryo production by enabling two rounds of SCNT from a single batch of oocyte collected through ovum-pick up or from abattoir ovaries.

Delaying nuclear maturation of the oocytes, from germinal vesicle to metaphase II, has been demonstrated using protein synthesis and kinase inhibitors such as cycloheximide and 6-dimethylaminoputine [7,8], phosphodiesterase inhibitors [9], cAMP modulators such as forskolin and 3-isobutyl-1-methylxanthine [10] with varied results on developmental competency. Alternatively, holding bovine oocytes at room temperature in simple holding media without additional drugs or inhibitors for 10 hours [11] or 16-18 hours [12] have similar embryo development rates as control oocytes that were not held. For equine oocytes, holding at room temperature overnight without inhibitors has been shown to have no negative consequences [13] and may even have increased embryo development rates [14]; and as a result, holding oocytes is presently a common practice in commercial equine embryo production. For SCNT, short-term holding of bovine oocytes for 6 hours with meiotic resumption inhibitors cilostamide resulted in reduced blastocyst development while natriuretic peptide type C supplementation during holding did not affect embryo development rates [15]. Longer holding of *Bos indicus* oocytes with butyrolactone I with brain-derived neurotrophic factor for 24 hours did not show detrimental effects on SCNT development rates [16]. It is not known if holding oocytes for an intermediate duration overnight without the addition of meiotic resumption inhibitors will have detrimental effects on development rates of handmade SCNT which requires more manipulation steps compared with micromanipulated SCNT.

The oocyte cellular environment has an extraordinary capacity to reprogram even terminally differentiated cells such as fibroblasts to become totipotent blastomeres within a time frame of normal early embryonic development, albeit at low efficiency. By day-7, SCNT blastocyst stage embryos have similar gene expression profile to in *in vivo* embryos produced by artificial insemination [17]. In comparison, epigenetic reprogramming of cells into induced pluripotent stem cells (iPSCs) using viral vectors takes weeks and is even less efficient than SCNT. While epigenetic reprogramming by SCNT can produce viable embryos, defects and incomplete epigenetic reprogramming can lead to higher rates of early embryo loss, fetal mortality, abnormal placentas, and fetal defects. Using pluripotent cells rather than terminally differentiated cells may reduce the degree of epigenetic reprogramming necessary to produce viable embryo by SCNT. In mice, embryonic stem cells and iPSCs donor cell SCNT embryos have higher developmental potential [18] compared with mouse embryonic fibroblast cell donor SCNT embryos [19]. Using umbilical cord mesenchymal stem cells resulted in SCNT embryos with closer gene expression profile to IVF embryos compared with SCNT embryos derived from fibroblast [20]. In horses, using bone marrow mesenchymal cells as somatic cell donors improved viability of SCNT produced foals compared with SCNT foals produced from fibroblasts, even though efficiency of embryo production was not different [21]. Therefore, it is possible that bovine iPSCs use as donor cell may improve SCNT efficiency and there may be benefits to offspring viability even if embryo production efficiency is not improved.

In this study, our objectives were: 1) to evaluate oocyte holding overnight in simple media on SCNT efficiency, enabling a second round of SCNT from a single oocyte collection date, and 2) compare the efficiency of bovine iPSCs as donor cells to conventional skin fibroblast on SCNT. We hypothesize that oocyte holding overnight can yield similar developmental outcomes after SCNT as oocytes processed on the day of aspiration.

## 2. Methods

### 2.1 Bovine fibroblast culture and cryopreservation

All reagents used for this experiment were purchased from Millipore Sigma (USA) unless stated otherwise. Bovine fibroblast cells were cultured and cryopreserved as previously described [5]. Fibroblasts were cultured in fibroblast culture media consisting of Dulbecco’s Modified Eagle Medium (DMEM, high glucose, GlutaMAX™ Supplement, pyruvate, Thermo Fisher Scientific, USA) with 10% Fetal bovine serum (FBS) and 1% penicillin-streptomycin (100X Solution, Thermo Fisher Scientific, USA) in a humidified incubator at 38.5°C an atmosphere supplemented with 5% CO_2_. Bovine fibroblast cryopreservation was accomplished using a slow-cooling protocol in straws. Briefly, fibroblast cells were detached from the plate by incubation with 0.25% trypsin-EDTA solution (Thermo Fisher Scientific, USA) for 5 minutes followed by the addition of culture media. The cell suspensions were centrifuged at 600 x g for 10 minutes and after discarding the supernatant, the pellet was diluted in cell freezing media (45% DMEM: 45% FBS: 10% dimethyl sulfoxide) to a concentration of 120,000 cells/mL, loaded into 0.25 mL straws. The straws were floated at 5 cm above the nitrogen vapor for 10 minutes then plunged to liquid nitrogen and stored.

Each experimental replicate was performed using cells expanded from warming a straw. Straws were removed from the liquid nitrogen tank and thawed in a water bath at 37°C for 30 seconds. Its content was added to a 15 mL tube containing 10 mL of warmed cell culture media and centrifuged for 10 minutes at 600 x g. Cells were plated in a 96-well plate in three wells with different concentrations four to five days before the somatic cell nuclear transfer procedure. At every replicate for both conventional and delayed matured oocytes, the well containing the cells with highest confluence was used to increase the proportion of cells in G0/G1 phase of the cell cycle. Cells were detached from the plates into suspension as described above for somatic cell nuclear transfer. For the reconstruction, only single cells of small size, round shape and intact membrane were selected.

### 2.2 Bovine induced pluripotent stem cell (biPSC) preparation

The methods for generation and sustenance of biPSCs have been previously described [22]. Bovine iPSCs were cultured over irradiated mouse embryonic feeders (iMEFs) [23] in GMTi medium [22]: DMEM/F12 containing N2 supplement, B-27 supplement, 1% non-essential amino acid supplement, 1% penicillin-streptomycin, 0.1 mM β-mercaptoethanol, 1.5 μM CHIR99021, 1 μM PD0325901, 0.5 μM A83–01, and 20 ng/ml hLIF or 20 ng/ml hIL6. with 10% FBS, 1% non-essential amino acids and penicillin-streptomycin. Propagation of the biPSCs were accomplished by dissociating colonies into single cells using TrypLETM (Thermo Fisher Scientific) and plating onto fresh iMEFs. Preparation of biPSCs for SCNT was similar with individual colonies manually picked then dissociated into single cells (**Fig 1**). All biPSC cultures were maintained in a humidified incubator at 37°C under an atmosphere of 5% CO_2_.

**Figure 1.**
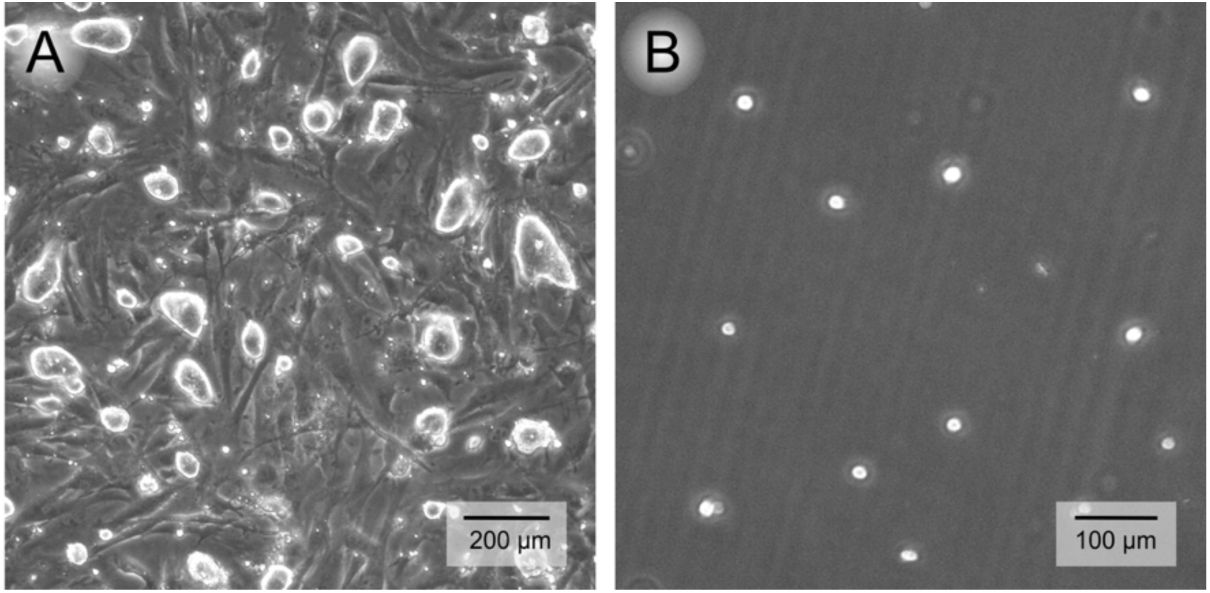
Panel A bovine induced pluripotent cell colonies plated over irradiated mouse embryonic fibroblast with characteristic colony morphology. Panel B is an images of single cell suspension after colonies were picked and dissociated to single cells for colony replating as well as preparation for somatic cell nuclear transfer.

### 2.3 Handmade Somatic Cell Nuclear Transfer

General bovine IVF media were from IVF Biosciences (United Kingdom). Oocyte manipulation media was based on Medium 199 (product number M2520) that contains Earle’s salts, L-glutamine, and 25 mM HEPES and supplemented with either 10% fetal bovine serum (M199FBS), or serum free with 0.01% polyvinyl alcohol (M199PVA).

Oocytes used this study were from bovine ovaries collected at a regional abattoir after transport for approximately 2 hours in normal saline at 23-25°C. Cumulus-oocyte complexes (COCs) were aspirated from follicles between 2-8 mm of diameter were aspirated with a vacuum pump (Cook Medical, USA). Grade 1 and 2 COCs were selected based on established classifications [24] and were assigned to one of two maturation timing treatment groups as shown in **Fig 2**.

**Figure 2.**
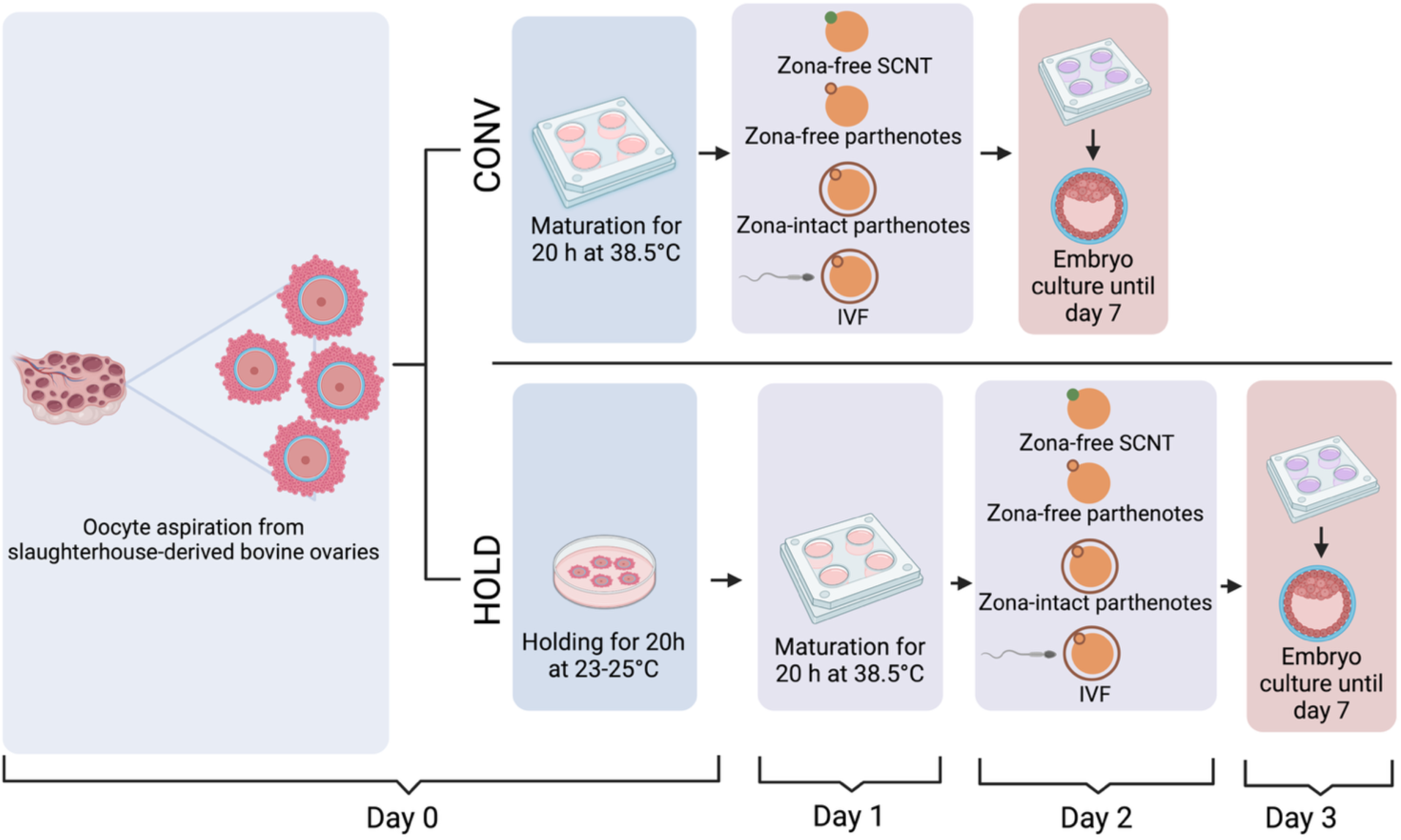
Experimental design: On Day 0, oocytes were aspirated from slaughterhouse ovaries and either placed in maturation for 20 hours (CONV group) or held at room temperature for the same time in holding media (HOLD). The next day (Day 1), the oocytes from the CONV group were divided into SCNT (using bovine fibroblast [bF] or bovine induced pluripotent stem cells [biPSC] as cell donors); IVF and zona intact and zona-free parthenogenetic activation groups as controls. At the same time, the HOLD group was placed in maturation for 20 hours. On day 2, the process from the previous day was repeated for HOLD group while CONV group oocytes were placed in culture. The same process was repeated in the following day for the HOLD group and oocytes were cultured for 7 days in total.

#### CONV Group

The conventional maturation (CONV) group were those where COCs were placed in maturation media for *in vitro* maturation immediately after selection. *In vitro* maturation was accomplished by culturing the COCs for 20 hours in four-well dishes containing 500 μL maturation medium (IVF Biosciences, United Kingdom) in an incubator at 38.5 °C, 5% CO_2_ and saturated humidity.

#### HOLD Group

The second group was the maturation holding group (HOLD) where COCs were placed in commercial holding media (ViGRO Holding Plus, Vetoquinol, USA) for 20 hours in 35mm dishes at room temperature in a humidified container. After the holding period ended, the COCs were moved to maturation media and incubated at 38.5 °C, 5% CO_2,_ and saturated humidity for another 20 h.

For each CONV and HOLD maturation timing treatment group, matured COCs were further assigned into one of four activation groups.

#### Zona Free SCNT activation

COCs in this group were denuded and zona pellucidae (ZP) removed, then used to produce handmade SCNT embryos using either bovine fibroblast or biPSC donor cells. Successfully fused embryos were parthenogenetically activated and cultured in microwells.

#### Zona Free parthenote activation

COCs in this group were denuded and ZP removed, then hemi-oocytes with nuclei were saved for parthenote activation together with the SCNT embryos to serve as SCNT processing control.

#### Zona Intact parthenote activation

COCs in this group were denuded to remove cumulus but were not subjected to ZP removal. The denuded oocytes were parthenogenetically activated with the zona-free groups to serve as parthenogenetic activation controls.

#### Zona Intact IVF activation

COCs in this group were left with intact cumulus and ZP then conventional *in vitro* fertilization was performed and served as *in vitro* embryo production control.

For the zona free groups and the Zona intact parthenote groups, the COCs were denuded to allow for assessment of maturation. Denudation was accomplished by placing the COCs in a 1.5 mL tube containing 100 µL of a solution of 0.1% of hyaluronidase in M199FBS. After incubation for 1 minute, the tube was vortexed (Fisherbrand Analog Vortex Mixer, Fisher Scientific, USA) for 30 seconds and the process was repeated three times. Denuded oocytes were then washed with M199FBS and removed from the tube for evaluation. Metaphase II oocytes were selected based on the visualization of the first polar body using a stereomicroscope.

The ZP was removed for the zona by treating the metaphase II oocytes with enzymatic digestion with 0.5% Pronase A in M199PVA, followed by extensive rinsing in M199FBS. Zona-free oocytes were incubated for 15 min in 5 μg/mL cytochalasin B in M199FBS, followed by manual bisection using a bisection blade (UltraSharp Splitting Blades, Shearer Precision Products, USA) under stereomicroscope.

After bisection, hemi-oocytes were stained with 10 μg/mL bisbenzimide (Hoechst 33342) diluted in M199FBS for 15 minutes. Hemi-oocytes were examined for the presence of a nucleus using an inverted epifluorescence microscope (Nikon TE 2000, USA) and the hemi-karyoplasts which are the hemi-oocytes with the nucleus were separated from the hemi-cytoplasts which were the hemi-oocytes without the presence of the metaphase II plate. The hemi-karyoplasts were kept in M199FBS at 38°C and used as Zona Free parthenote activation group.

For the Zona Free SCNT activation group, embryo reconstruction was performed by a brief exposure of two cytoplasts to 500 μg/mL phytohemagglutinin in M199PVA, followed by adhesion of a somatic cell. First, one hemi-cytoplast was moved to a single somatic cell and the pair was then moved to another hemi-cytoplast in a way that the cell stayed in between the two oocyte halves (**Fig 3**). Reconstructed structures were briefly rinsed in electrofusion (EF) medium (0.3 M mannitol, 0.05 mM CaCl_2_ 2H_2_O, 0.1 mM MgSO_4_ 7H_2_O, 0.5 mM HEPES, 0.01% PVA) and moved to a 32-mm gap fusion chamber (BTX 453, Harvard Apparatus, USA) containing 650 μL EF medium coupled to an electrofusion apparatus (BTX 830 Electro Square Porator, Harvard Apparatus, USA). Reconstructed structures were electrofused by one pulse of 1.2-kV/cm for 40 μs, and they were cultured individually in 5 μL micro drops of M199FBS under mineral oil, at 38.5 °C, on a warm plate, for 45–60 min until fusion evaluation (Supplemental video S1).

**Figure 3.**
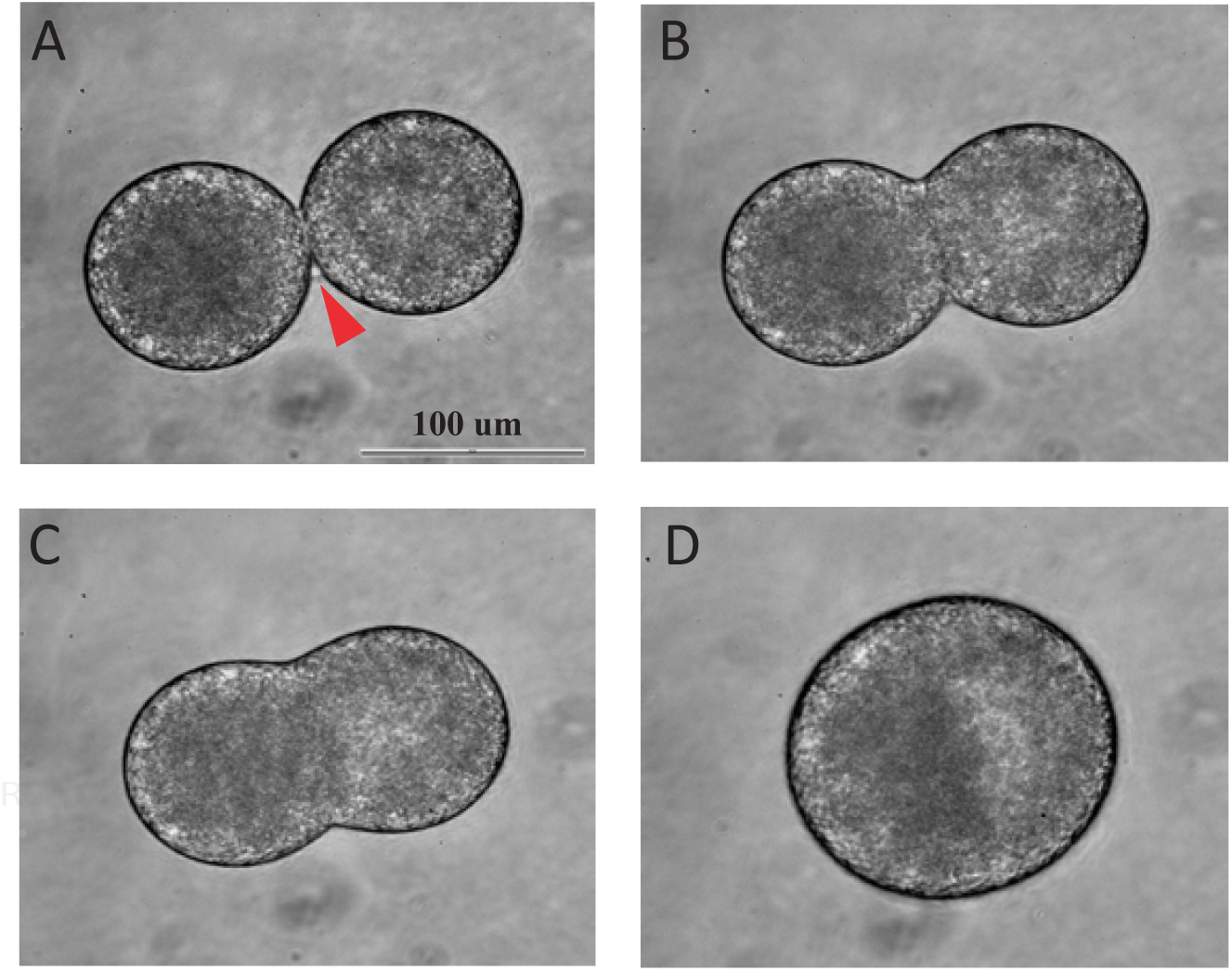
Time series of embryo reconstruction and fusion. Panel A shows a reconstructed embryo consisting of two hemi-cytoplasts and a donor cell (arrow), Panel B shows the donor fibroblast fusing with the hemi-cytoplasts that are in the early stages of fusion, Panels C and D shows the fusing of hemi-cytoplasts to form an embryo.

### 2.4 Oocyte activation and Embryo Culture

Parthenote activation was performed on the successfully fused cloned embryos in the Zona free SCNT activation group, the Zona free parthenote activation group which were the karyoplasts pairs, and the Zona-intact parthenote activation group. The oocytes were chemically activated in 5 μM ionomycin diluted in M199FBS for 5 min, followed by individual incubation in 5 μL microdroplets with 2-mM 6-DMAP diluted in *in vitro* culture media (IVC, IVF Biosciences), under mineral oil, for 4 h. The fusion-activation interval was approximately 2 hours, and the oocytes were activated under 30 hours after the onset of maturation.

For the Zona intact IVF activation group, conventional *in vitro* fertilization was performed on the same batch of oocytes used fr the SCNT experiment, using cryopreserved semen from a fertility-proven bull. The straw was warmed by immersing the straw in a water bath at 37° C for 20 seconds. Spermatozoa was deposited in a 15 mL conical tube containing 5 mL of sperm washing media (BO-Semen Prep, IVF Biosciences, United Kingdom), washed by centrifugation at 350 x g for 5 minutes. After removing the supernatant, the pellet was diluted with 1 ml of sperm washing media (BO-Semen Prep, IVF Biosciences, United Kingdom) and centrifuged again for 5 minutes at 350 x g. The semen concentration was assessed in an automatic cell counter (Nucleocounter SP-100, ChemoMetec, Denmark) and 1 × 10^6^ sperm cells/mL concentration was adjusted and added proportionally to the wells containing the oocytes. The matured COCs were washed three times in 100 μL droplets of BO-IVF (IVF Biosciences, United Kingdom) then transferred to a 5-well plate containing 500 μL of BO-IVF (IVF Biosciences, 61003) per well overlaid with mineral oil. Fertilization was carried out for 18 hours at 38.5° C in a humidified atmosphere of 5% CO_2_ in air. After fertilization, presumptive zygotes were removed from the fertilization plates and denuded by vortexing (Fisherbrand Analog Vortex Mixer, Fisher Scientific, USA), using the same method described above for the oocytes used for nuclear transfer.

Embryos were cultured for 7 days in embryo culture media (BO-IVC, IVF Biosciences, United Kingdom), overlaid with mineral oil in humidified atmosphere of 5% O_2_, 5% CO_2_, and 90% N_2_ at 38.5°C. Zona free SCNT embryos were cultured individually in a Well of the Well system (WOW) [25] made with a special needle (BLS Limited, Hungary) within the culture media droplet, or in a commercial WOW time lapse plate (Primovision, Hungary) to monitor embryo development as shown in **Fig 4** (Supplemental video S2). A total of 10-15 cloned embryos were cultured in 50 μL droplets of IVC media under oil. Zona free parthenote controls were also cultured in WOW; however, two hemi-karyoplasts were cultured per microwell to compensate the difference in total embryo volume compared with SCNT fused embryos. Zona intact parthenote and Zona intact IVF controls were cultured in 5-well dishes (Vitrolife, Sweden) containing 500 μL of culture media (BO-IVC, IVF Biosciences, United Kingdom), overlaid with mineral oil. A total of 50 embryos were cultured per well.

**Figure 4.**
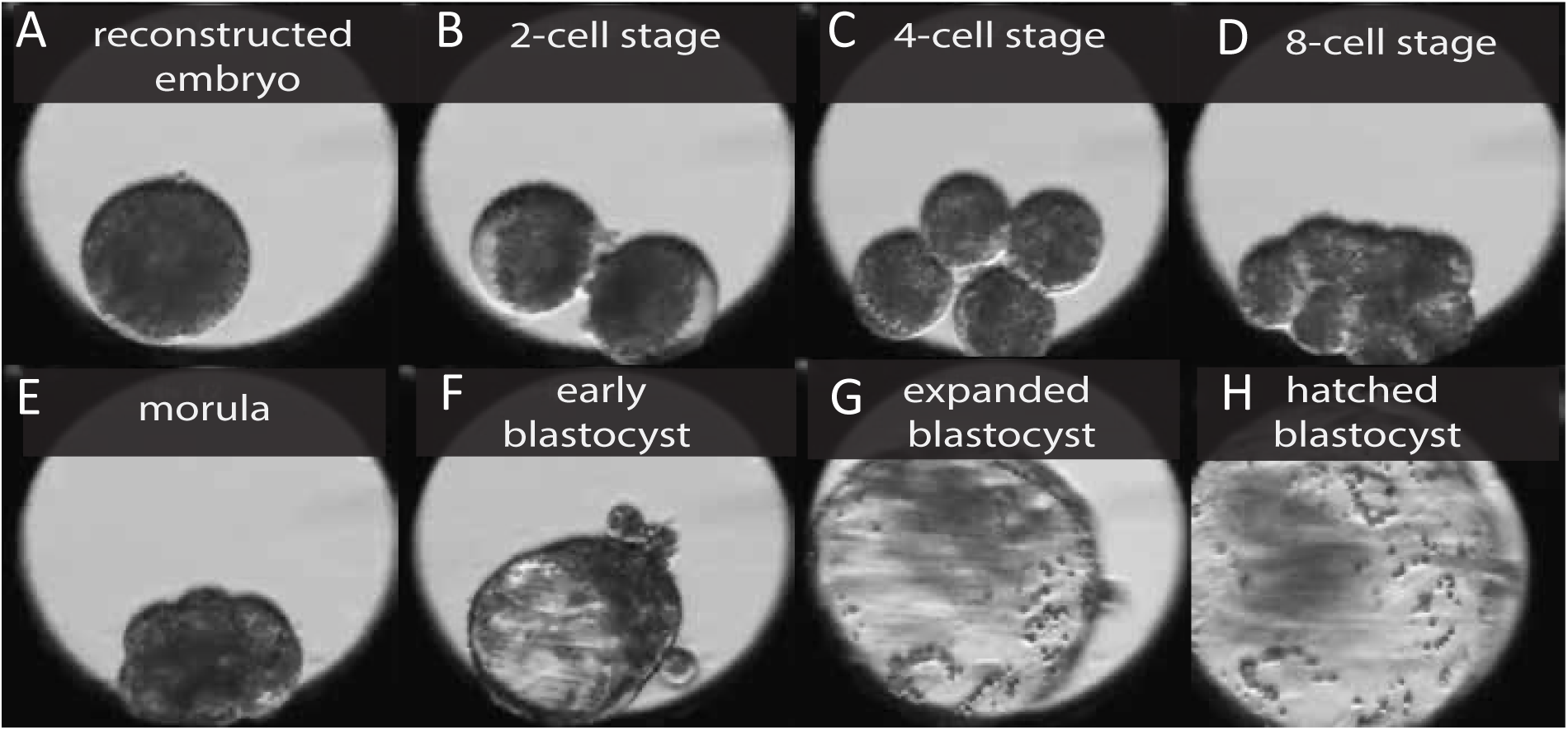
Time-lapse monitoring of handmade clone embryo culture for seven days following activation (A-H). The handmade clone embryos are zona-free and requires a confined space in a WOW culture system.

Embryos cleavage rate assessment was performed 48 hours (Day 2) after fertilization or chemical activation (set as time = 0 hour). Development to the blastocyst stage was evaluated on Day 7, with blastocysts classified morphologically according to stage of development (Stages 5–9) and morphological quality (Grades 1 to 3), adapted from the guidelines of the International Embryo Technology Society (IETS) for *in vivo*-derived embryos.

### 2.5 Statistical analysis

All analyses were performed using JMP Pro version 17 (SAS Institute, Cary, NC, USA) and R (www.R-project.org). Because of biological and technical variability inherent to oocyte manipulation and blastocyst formation, unpaired analyses were used. Categorical outcomes representing binary developmental success were analyzed by logistic regression, and interval-level outcomes were analyzed by linear regression ANOVA to determine differences between groups. For logistic models, odds ratios with 95% confidence intervals were calculated to assess relative likelihoods of developmental success between donor cell types. All results are reported as mean ± standard error, and significance was set at P ≤ 0.05.

## 3. Results

A total of 1,823 oocytes were used in this experiment (1,372 for zona free SCNT, 185 as zona intact parthenote controls and 266 for zona intact IVF controls. For the zona free parthenote controls, 220 pairs of hemi-karyoplasts were cultured. Data from 6 replicates (for all groups except zona intact IVF and biPSC, which had 3 replicates each performed) were used for the analyses. Recovery after stripping (CONV = 92.4±4.1, HOLD = 91.7±2.4), maturation rates (CONV = 70.2±3.2, HOLD = 67.2±4.0) and survival rates either after zona removal (CONV = 98.3±0.5, HOLD = 98.9±0.6) or oocyte bisection (CONV = 81.4±4.0, HOLD = 81.13.0) did not differ between groups (*P* > 0.05). In addition, the proportion of oocytes selected for reconstruction was also not significantly different (CONV = 50.1±1.3, HOLD = 51.1±2.2) as shown in **Table 1**.

**Table 1.** Comparison between oocytes *in vitro* matured for 20 hours (CONV) vs. oocytes held for 20 hours at room temperature followed by another 20 hours of maturation regarding their recovery after maturation (HOLD), maturation rate, survival after zona removal and bisection, in addition to the number of hemi-oocytes selected for nuclear transfer.

| Group | Oocytes in maturation | Oocytes recovered (n, %) | Matured oocytes (n, %) | Zona removal (n) | Zona removal (n, % survival) | Oocyte bisection (% survival) | Hemi oocytes selected (%) |
| --- | --- | --- | --- | --- | --- | --- | --- |
| CONV | 692 | 640, (92.5%) | 486, (70.2%) | 351 | 345, (98.3%) | 286, (81.4%) | 179, (50.1%) |
| HOLD | 680 | 624, (91.8%) | 457, (67.2%) | 351 | 347, (98.9%) | 285, (81.2%) | 179, (50.1%) |

There were no significant differences in these parameters between groups.

Fusion rate was not significantly different between CONV (42.6±6.0) and HOLD (50.3±9.8) groups when bovine fibroblasts were used (**Table 2**). However, for the biPSC, oocytes matured conventionally showed a higher fusion rate (85.7±8.2) compared with HOLD group (50.5±8.8) (*P* < 0.05). Since all non-fused oocytes were re-fused, the total number of embryos cultured was similar for the biPSC cells (CONV = 18/114, 15.8%, HOLD = 22/104, 21.1%). This is helped by the high refusion rate for the HOLD group (91.7±8.3) compared with the CONV group (16.7±16.7) (*P* < 0.05). The percentages do not result in 100% because the refusion rates are calculated based on the number of non-fused embryos and not the total. When comparing fusion success rates between donor cell types, the proportion of embryos that successfully fused and cultures was higher (Odd ratio 5.9, 95% C.I. 1.8 – 19.8) in the biPSC (93.0%) compared with bovine fibroblasts (69.3%) (*P* < 0.05).

**Table 2.** Comparison of fusion and re-fusion rates of two enucleated hemi-oocytes with bovine fibroblast or a bovine-induced pluripotent stem cell (biPSC) between conventional maturation (CONV) or maturation after holding oocytes at room temperature (HOLD).

| Group | Bovine fibroblast |  |  |  | biPSC |  |  |  |
| --- | --- | --- | --- | --- | --- | --- | --- | --- |
|  | Cytoplasmic couplets | Fused (%) | Re-fused (%) | Cultured (n, %) | Cytoplasmic couplets | Fused (%) | Re-fused (%) | Cultured (n, %) |
| CONV | 114 | 42.6±6.0 | 52.5±19.2 | 79, 69.3% | 20 | 85.7±8.2<sup>a</sup> | 16.7±16.7<sup>a</sup> | 18, 90.0% |
| HOLD | 104 | 50.3±9.8 | 65.6±13.8 | 72, 69.2% | 23 | 50.5±8.8<sup>b</sup> | 91.7±8.3<sup>b</sup> | 22, 95.7% |
| Combined | 218 |  |  | 151, 69.3%<sup>c</sup> | 43 |  |  | 40, 93.0%<sup>d</sup> |

<sup>a,b</sup> Different superscripts within columns indicate differences ( $P < 0.05$ ) between CONV and HOLD groups
<sup>c,d</sup> Different superscripts within rows indicate differences ( $P < 0.05$ ) between bovine fibroblast and biPSC cells

There was no effect of maturation group on cleavage in the zona intact IVF (CONV = 82.8±2.9, HOLD = 79.0±3.5), or zona intact parthenote controls (CONV = 87.6±2.6, HOLD = 87.8±2.6) and in the SCNT groups with both fibroblast (CONV = 88.3±10.4, HOLD = 86.0±8.1) and biPSC (CONV = 94.4±5.5, HOLD = 87.5±12.5) cells. There was a significant increase in the cleavage rate of the zona-free parthenote in the HOLD group (97.2±1.3) compared with the CONV group (91.6±4.6) (*P* < 0.05).

Maturation treatment had no significant effect on the blastocyst rates, except for the zona intact parthenotes, where the CONV group (55.9±5.6) showed a higher blastocyst rate compared with the HOLD group (39.6±5.1) (*P* < 0.05). Although differences were not significant, CONV group oocytes tended to yield higher blastocyst rates for both cell types in SCNT embryos. Among donor cell types, blastocyst formation rates of reconstructed embryos using biPSCs were consistently higher than those using fibroblast donor cells in both maturation groups (CONV = 33.3 ± 16.7 vs 21.9 ± 12.6, P < 0.05; HOLD = 26.4 ± 6.0 vs 14.7 ± 8.9, P < 0.05; Table 3).

**Table 3.** Cleavage and blastocyst rates of oocytes *in vitro* matured for 20 hours (CONV) or held for 20 hours at room temperature before another 20 hours of *in vitro* maturation. Data for the two different lines of donor cells fibroblast (SCNT-fibroblast) or biPSCs (SCNT-biPSC) and technical controls namely standard *in vitro* fertilized (IVF) embryos, zona-intact and zona-free parthenogenetically-activated embryos are shown

| Parameter | Group | IVF | Zona-intact<br>parthenotes | Zone-free<br>parthenotes | SCNT-fibroblast | SCNT-biPSC |
| --- | --- | --- | --- | --- | --- | --- |
| Cleavage (%) | CONV | 82.8±2.9 | 87.6±2.6 | 91.6±4.6<sup>a</sup> | 88.3±10.4 | 94.4±5.5 |
|  | HOLD | 79.0±3.5 | 87.8±2.6 | 97.2±1.3<sup>b</sup> | 86.0±8.1 | 87.5±12.5 |
| Blastocyst (%) | CONV | 31.8±5.5 | 55.9±5.6 | 38.6±8.4 | 21.9±12.6<sup>c</sup> | 33.3±16.7<sup>d</sup> |
|  | HOLD | 27.6±5.6 | 39.6±5.1 | 40.5±3.6 | 14.7±8.9<sup>c</sup> | 26.4±6.0<sup>d</sup> |

<sup>a,b</sup> Different superscripts indicate differences ( $P < 0.05$ ) between CONV and HOLD groups; <sup>c,d</sup> Different superscripts indicate differences ( $P < 0.05$ ) between fibroblast and biPSC.

When considering the total efficiency of embryo production after reconstruction, there was a significant effect of cell type on fusion rates. Combined with numerical differences in cleavage and blastocyst rates between bovine fibroblast and biPSC, the overall cleavage and blastocyst rates out of total reconstructed embryos was significantly higher in biPSC compared with bovine fibroblast for cleavage rates (Odds Ratio = 3.4, 95% C.I. 1.5 – 8.1, P = 0.0047) and blastocyst rates (Odds ratio 2.7, 95% C.I. 1.2 – 5.9, P = 0.0128).

## 4. Discussion

Cloning by somatic cell nuclear transfer had been used to replicate high value animals, in conservation to rescue endangered and even extinct species, to generate transgenic animals among other uses. After more than 25 years of the first report of the technique in cattle, it is still a time-consuming procedure that requires high-skilled personnel and expensive equipment to be performed. The development of the handmade cloning technique, allowed to produce embryos without micromanipulators, however this variation and other adaptations require the same or more time and skills in order to be successful. As an example, in this experiment for approximately 120 oocytes, 30 minutes were used for polar body selection, 30 minutes for zona removal, 60 minutes for bisection and recovery, another 60 minutes for selection of cytoplasts in addition to another 60 minutes for reconstruction of the pairs, cell preparation and fusion to the hemi-oocytes. Including another two hours between fusion and activation, the total time spend manipulating the oocytes exceeds 6 hours after the end of maturation.

As efficiency increases depending on the skill and there are variations between individuals and the technique used to produce SCNT embryos, our objectives were to maximize the use of oocytes in a single collection, performing the SCNT technique over two consecutive days. According to previous reports [13], it is possible to produce blastocysts at the same rate with oocytes held at room temperature, however there is no information about SCNT embryos using the same approach. One of our concerns was that the addition of aged oocytes and the extended manipulation and exposure to different chemicals would negatively affect the embryo production. Even though it was showed that aged oocytes can support blastomere cloning, and that electrofusion is enough to induce oocyte activation [26], we wanted to compare the efficiency of the exact same protocol in both days of SCNT. Starting with maturation, no significant differences were seen between HOLD and CONV groups. Since only visual assessment performed, we have no information about the number of degenerated oocytes or arrest in other stages. However, the recovery after stripping was not different between groups (> 90%) what suggests that the oocytes held in the HOLD group had similar quality than the conventionally matured. This is corroborated by the fact that survival after chemical removal of the zona pellucida and manual bisection were similar between groups.

Using handmade cloning it is possible to remove a small amount of cytoplasm around the polar body and use the remaining oocyte as a recipient. In our experiment, on the other hand we aimed to bisect the oocytes in two equal parts, which reflects that approximately 50% of the halves were selected as recipients, regardless of the experimental group. It is also important to stress that number of oocytes used for SCNT was also reduced since part of the total was used as zona free or zona-intact controls.

Fusion rates can vary depending on the cell type, passage and methods used for fusion. In our case, neonatal fibroblasts presented a fusion below most reports in both CONV and HOLD groups, but since all non-fused pairs were re-fused, approximately 69% of the total number of reconstructed couplets were fused. Due to the low number of embryos for culture, they were activated and cultured together, making impossible to differentiate what were the development rates for each group. For the biPSC, the fusion rates were significantly higher for the CONV group, similar to what was previously reported for this type of cell [27]. However, the re-fusion rates were much lower for the same group while higher rates were seen in the HOLD group, suggesting that in our conditions, for biPSCs, the parameters for fusion were not adequate or these cells are more resistant to fusion. The overall fusion rates for these cells was approximately 93% which was comparable to previous reports [27,28].

Cleavage rates were similar between CONV and HOLD groups for embryos produced by IVF, parthenogenesis and SCNT. Interestingly, the rates for the zona-free parthenogenetically activated group were higher in the HOLD than the CONV maturation group. A possible explanation is that it is more difficult to differentiate cleavage from fragmentation within the WOW system. Aged oocytes are more prone to fragmentation [29] and that might be the one of the reasons the HOLD group presented higher cleavage rates. In addition, the zona-free parthenotes were the only group in the experiment where the two hemi-oocytes were cultured by aggregation. That could lead to one of the halves having better quality and dividing while the other half is arrested, allowing an aged oocyte more changes to divide. According to Choi et al., [12] oocytes held for 18 hours plus 18 hours of maturation (total 36 hours) also had similar cleavage rates than the conventionally matured oocytes when used for IVF. In our experiment the oocytes were held for 2 hours longer, and our rates were still similar. When they tested oocytes matured for longer, up to 40-42 hours (18 hours holding plus 22-24 of maturation) the cleavage rate decreased but, in our conditions, oocytes used for IVF, parthenogenesis or SCNT still can cleave at the same rates of conventionally matured oocytes. In addition, in our conditions, the use of oocytes 40 hours after initial culture produced the same cleavage rates than conventional maturation. However, our maturation time was shorter, which might suggest that holding time can be increased but once maturation is started, the oocytes have to be fertilized or activated in a timely manner.

Blastocyst rates after *in vitro* fertilization, in our experiment, did not differ between the HOLD and CONV groups, similarly to what was previously reported [11,12]. *In vitro* fertilization requires sperm capacitation and it peaks around 10-12 hours after the start of co-incubation with the oocyte with sperm [30]. Oocytes also acquire their highest competence around 30 hours after the onset of maturation [31], which coincides with the fertilization time. Similarly, one-cell embryos produced by SCNT or parthenogenesis in our conditions, were activated before the 30 hours mark. However, activation at 27 hours with fusion at 24 hours achieved high blastocyst rates in previous reports), more than two times the rate found in our conditions. Interestingly, when oocytes were held for 20 hours and fertilized or activated the same way as their conventional counterparts, in all experiment groups, the blastocyst rates did not significantly decrease. Even though we did not test how long the maturation can be delayed, we showed that up to 20 hours of holding at room temperature do not affect embryonic development, which is a longer time than previously reported. In addition, the blastocyst rates for the HOLD group were not different for SCNT embryos of both cell lines, suggesting that electrical activation did not occur because the oocytes were not truly aged, or the development was not affected after a subsequent chemical activation. Moreover, the blastocyst rates achieved in our experiment for fibroblasts and pluripotent stem cells were similar to previous reports for these type of cells [27,32].

Overall embryo production efficiency was higher in biPSC compared with bovine fibroblast driven in part by the higher overall fusion rates. Consistent with this, embryos reconstructed with biPSCs showed superior developmental outcomes, with blastocyst formation rates of 33.3% compared with 21.9% for fibroblast donors. This pattern was observed in both maturation treatments, indicating that oocyte holding was non-detrimental and that the advantage conferred by pluripotent donor cells was independent. When considering all reconstructed embryos, biPSCs also yielded higher overall cleavage and blastocyst development, reflected by logistic regression-derived odds ratios 3.4 and 2.7, respectively. Although the basis for the higher fusion rate observed with biPSCs remains unclear, these findings suggest that nuclear transfer using pluripotent donor cells can enhance developmental potential at multiple stages, likely by reducing the degree of epigenetic reprogramming required after SCNT. This concept is supported by prior evidence in cattle where SCNT using pluripotent donor blastomeres (8-64 cell stage embryos) achieved substantially higher cloning efficiency than with somatic cells [33].

In conclusion, delaying the maturation for 20 hours at room temperature in a simple media allows oocytes not only to develop into IVF blastocysts but also to SCNT embryos with different cell lines. This shows for the first time that it is possible to produce embryos by handmade cloning in two consecutive days using oocytes collected in a single day, reducing costs and time to process the ovaries or to perform ovum pick up. Moreover, biPSC donor cells yielded higher blastocyst rates, suggesting improved nuclear reprogramming during SCNT. Together, these findings confirm our hypothesis that oocytes can be held for SCNT without compromising competence and that pluripotent donor cells can increase cloning efficiency. The ability to hold oocytes before maturation, as routinely done in the equine industry, could also provide greater flexibility for the commercial laboratories, allowing maturation to begin at convenient times for resources and personnel. This would also enable shipment of viable oocytes from remote collection sites for *in vitro* embryo production.

## Supplemental Material

Video S1: Time lapse video of the fusion of a reconstructed embryo consisting of two hemi-cytoplasts derived from manual bisection of bovine oocytes and a neonatal fibroblast cell over 60 minutes following electrofusion.

Video S2: Time lapse video of a handmade clone embryo culture for seven days. This handmade clone embryo was produced by fusing hemi-cytoplasts with a bovine induced pluripotent cell.

